# Chronic administration of the hydrogen sulfide prodrug SG1002 partially protects against erectile dysfunction resulting from long-term androgen deprivation

**DOI:** 10.1101/2025.05.26.656242

**Authors:** Colin M. Ihrig, Clifford J. Pierre, Tooyib A. Azeez, Justin D. La Favor

## Abstract

**Aims:** Androgen deprivation therapy is a common treatment strategy for prostate cancer, although erectile dysfunction (ED) often coincides as an undesirable side-effect. Hydrogen sulfide (H_2_S) is an endogenous gasotransmitter with vasodilatory, anti-inflammatory, and antioxidant-like properties. H_2_S therapies are being developed for cardiovascular disease management, although the properties of H_2_S may also protect the erectile system.

**Materials and methods:** 14-week-old male C57Bl/6 mice were subjected to sham surgery or castration, with castrated mice remaining untreated or treated orally with low- or high-doses of the H_2_S prodrug SG1002 over the five-week intervention. Erectile function was assessed by intracavernous pressure and mean arterial pressure during cavernous nerve stimulation. Vascular reactivity of the corpus cavernosum (CC), internal pudendal artery (IPA), and internal iliac artery (IIA) were assessed by dose-dependent responses to vasodilatory, vasocontractile, and neurogenic stimuli in myograph systems. CC contents of proteins related to cellular autophagy, antioxidant defense, and mitochondrial dynamics were assessed by immunoblotting. Fibrotic remodeling was assessed by Massons trichrome staining.

**Key findings:** Both doses of SG1002 provided an equivalent and moderate protection on erectile function against long-term androgen deprivation. Castration-induced alterations of several mechanisms of vasodilation and vasoconstriction of the CC and IPA were substantial, while alterations of the IIA modest, with subtle effects of SG1002 treatment across the vascular beds. SG1002 partially protected against castration-induced fibrotic remodeling of the IPA.

**Significance:** H_2_S therapy provides a modest but potentially clinically relevant protection of erectile function and health of the erectile structures against the harshly damaging effects of chronic androgen deprivation.

## 1. Introduction

Prostate cancer is one of the most commonly diagnosed cancers globally with an estimated 1.4 million new cases in 2020 [1]. Men undergoing treatment for prostate cancer often will experience hypogonadism as part of their standard treatment to slow and/or shrink prostate cancer growth. This is achieved via a form of chemical castration called androgen deprivation therapy (ADT) through the utilization of gonadotropin-releasing hormone (GnRH) agonists, antagonists and antiandrogens [2]. Hypogonadism resulting from the use of ADT is associated with increased rates of erectile dysfunction (ED) and increased exposure of the erectile tissue to a hypoxic environment resulting in a pro-fibrotic environment [3,4]. ED is defined as the inability to achieve or maintain an erection capable of satisfactory sexual intercourse. Individuals undergoing ADT experience decreased sexual functioning which is affected by the duration of therapy [3]. To mimic the effects of chemical castration on vascular and erectile function that these patients are subjected to, double orchiectomy animal models are commonly utilized. Such rodent models have demonstrated significant deterioration in vascular function of pre-penile arteries, as well as the corpus cavernosum [5–7].

Hydrogen sulfide (H_2_S) is an endogenous gasotransmitter that has become a major focus in vascular regulation. H_2_S has been shown to work directly with another prominent gaseous signaling molecule nitric oxide (NO) as a vascular tone regulator [8,9]. H_2_S also acts directly as a vasodilator via hyperpolarization of the underlying smooth muscle. H_2_S production occurs via the breakdown of cysteine by three main enzymes cystathionine γ-lyase (CSE), cystathionine-β-synthase (CBS), and 3-mercaptopyruvate sulfotransferase (3MST). Additionally, H_2_S acts in an antioxidative capacity, partly through stimulation of Nrf2 transcriptional activity [10]. In conjunction with the anti-inflammatory effects, H_2_S has also shown been shown to blunt fibrotic development through its effects on vascular smooth muscle cells [11]. Castration has been demonstrated to significantly increase corpus cavernosum (CC) fibrosis of rats [12,13]. One of the potential pathways attributed to this development in fibrosis is inhibition of autophagy and promotion of apoptosis of the CC smooth muscle cells [12]. In addition to its effects on autophagy, H_2_S has also been found to positively maintain balance over mitophagy and mitochondrial dynamics of fusion and fission [14]. Sun et al. found H_2_S supplementation to stimulate pathologically impaired mitophagy via increased translocation of parkin into the mitochondria in cardiac muscle in a db/db mouse model [15].

With vascular remodeling and fibrosis being closely tied to the development and progression of ED, decreasing the inflammatory burden on the system may prove beneficial to both vascular and erectile health and function. Testosterone has been demonstrated to have a stimulatory effect on H_2_S induced vasodilation in rat aortas [16]. Additionally, Cui et al. found that administration of testosterone was able to restore erectile function and blunt the increased fibrotic molecular changes associated with castration in rats [13]. Testosterone supplementation also has positive effects on erectile health and function [17]. However, testosterone supplementation in individuals with prostate cancer is counterproductive to the suppression of androgen production and binding by most aforementioned frontline therapeutic strategies.

Importantly, stimulation of H_2_S induced vascular homeostasis in models absent testosterone may prove to have a protective effect on the vasculature and erectile tissue and thus improve long-term patient outcomes. Previous work demonstrates that an orally active H_2_S prodrug SG1002 is able to augment circulating H_2_S levels and have positive cardiac and vascular health outcomes in both mice and humans under various conditions associated with cardiovascular disease [18–23].

Our lab has demonstrated that castration induces the downregulation of several antioxidant genes in the mouse CC, including Nqo1, Sod2, Sod3, Gclc, Gpx1, Prdx3, and Prdx5. This downregulation reversible in all but Sod3 with high doses of the antioxidative compounds resveratrol and MitoQ [5]. Utilizing a western diet model we have also demonstrated the ability of 20 mg/kg/day of SG1002 to increase CC levels of several antioxidant proteins [24]. Therefore, the aim of this study was to determine if SG1002 treatment protects against the detrimental effects of androgen deprivation on erectile health. We assessed erectile function in the intact animal, as well as the vascular physiology of the CC and the pre-penile arteries involved in the erectile response. We assessed these tissues for fibrotic remodeling, while we also evaluated the influence of SG1002 treatment on proteins related to autophagy, mitochondrial dynamics, and cellular antioxidant defense in the end organ CC. We investigated a low and a high dose of SG1002 to determine potential dose-dependency of any observed effects.

## 2. Methods

### 2.1 Experimental Animals and Study Design

136 Male C57Bl/6J mice received from Jackson Laboratories (Bar Harbor, ME, USA) were divided into four cohorts, one cohort that underwent sham castration surgery (Sham) and three cohorts that underwent castration. Castrations were performed at 14 weeks of age under the administration of anesthesia via isoflurane. Animals received subcutaneous injections of buprenorphine post-surgery and again 5 hours post-surgery at a dosage of 0.05 mg/kg body weight. The sham group and one castrated cohort (Cast) remained on a standard chow diet (5001 PMI Laboratory Rodent Diet, Inotiv, Madison, WI, USA). The remaining two cohorts were placed on identical diets three days prior to castration that were supplemented with SG1002 (Sulfagenix, Cleveland, OH, USA) at concentrations designed to deliver approximately 20 mg/kg/day (CLS) or 100 mg/kg/day (CHS) that were administered throughout the experimental period. Functional assessment and experimentation occurred 5 weeks post castration to mimic an extended period of androgen deprivation. All experimental procedures were approved by the Animal Care and Use Committee of Florida State University (protocol 202100052).

### 2.2 Erectile Function Assessment

Erectile function was assessed through measurement of intracavernosal pressure (ICP) and mean arterial pressure (MAP) as described previously [25]. Briefly, mice were anesthetized with an intraperitoneal injection of 90 mg/kg ketamine and 10 mg/kg xylazine. The left carotid artery and left crus were cannulated with polyethylene tubing filled with 100 U/ml of heparinized saline connected to pressure transducers (ADInstruments, Sydney, Australia), allowing for the continuous measurement of mean arterial pressure (MAP) and intracavernous pressure (ICP) through connection to a PowerLab data acquisition system (ADInstruments) via a bridge amplifier (#FE221, ADInstruments) with LabChart Pro software (ADInstruments). The cavernous nerve was stimulated with bipolar electrodes via a square pulse stimulator (Grass Instruments, Quincy, MA, USA) at a frequency of 10 Hz with a 5 ms pulse width for 60 s stimulation periods, separately at 0.5, 1, 2, and 4 V of stimulation. Mice were sacrificed 10 minutes following the final stimulation by double thoracotomy and exsanguination of the vena cava. Erectile function was assessed by the peak ICP-to-MAP ratio and the area-under-the-curve (AUC) of the ICP tracing during stimulation normalized to MAP (AUC/MAP), which represent the ability to achieve and maintain an erection, respectively.

### 2.3 Ex vivo Vascular Reactivity of the Corpus Cavernosum and Pre-Penile Arteries

In a separate set of mice from the erectile function assessment, mice were deeply anesthetized with an intraperitoneal injection of 90 mg/kg ketamine and 10 mg/kg xylazine and sacrificed by double thoracotomy and exsanguination of the vena cava. Penile tissue was removed under a dissection microscope, and the penile shaft was separated from the glans penis. The penile shafts were placed in ice-cold Krebs solution of the following composition (in mM): NaCl 130, KCl 4.7, KH2PO4 1.18, MgSO4 1.18, NaHCO3 14.9, dextrose 5.6, CaCl2 1.56, and EDTA 0.03 dissolved in distilled water. All components of the Krebs solution were purchased from Sigma Aldrich (St. Louis, MO, USA). The corpus spongiosum, dorsal vein, and connective tissues were carefully excised from the penile shaft in chilled Kreb’s solution. Two segments of CC were obtained via cutting the CC along the septum splitting into two equal strips. CC strips were mounted in a DMT 820MS muscle strip myograph system (Danish MyoTechnology (DMT), Aarhus, Denmark) for isometric tension measurement and recording with LabChart software (ADInsruments) [5]. Two segments (1.5-2.0 mm length) internal iliac artery (IIA) and two distal segments of the internal pudendal arteries (IPA) were dissected from each animal and mounted in a DMT 620M wire myograph system using 25 µm tungsten wire as previously described [26]. Myograph systems were coupled to a PowerLab 8/30 data acquisition system (AD Instruments, Dunedin, New Zealand) for continuous recording of isometric tension measurement with LabChart Pro software (AD Instruments). Tissues were bathed in Kreb’s solution maintained at 37°C and continuously aerated with a 95% O2 and 5% CO2 mixture.

Tissues were allowed to equilibrate for 1 h, after which the length-tension relationship of the IPA and IIA segments were tested by gradually increasing the diameter until the transmural pressure exceeded 100 mmHg, and the vessel diameters were set to 90% of the internal circumference that elicited 100 mmHg transmural pressure calculated by DMT Normalization Software (LabChartPro, AD Instruments), while CC strips were stretched to a resting tension of 4 mN. Tissues then equilibrated for 45 min, then tissue viability was tested with high potassium (120 mM) Krebs solution, with KCl substituted for NaCl. Tissues were washed successively every 10 min for at least 30 min or until tissues returned to their stable baseline resting tension between the following dose-response challenges.

Agonist-mediated vasoconstriction was tested by cumulative dose-response applications of 0.001 µM-10 µM of the α_1_-adrenoreceptor agonist phenylephrine (PE), 0.3 nM -100 nM (II and IPA) or 0.001 µM-3 µM (CC) of the thromboxane A_2_ receptor agonist U-46619, 1 nM-100 nM endothelin-1 (ET-1), and electrical field stimulation (EFS). EFS was applied with a CS8 stimulator (DMT) controlled by DMT MyoPULSE software. The following stimulation parameters for EFS were used: frequencies of 0.5-32Hz (IIA and IPA) or 1-32Hz (CC), 10 s stimulation trains, 2 millisecond pulse width at 20 V (IIA and IPA) or 30 V (CC), and 2 minutes between each stimulation train. Vasodilatory capacity was tested following 1 µM PE (IIA and IPA) or 10 µM PE (CC) pre-constriction by cumulative dose-response applications of 0.001 µM-10 µM acetylcholine (ACh), 0.001 µM-3 µM of the nitric oxide donor sodium nitroprusside (SNP), 10 µM-300 µM of the rapid releasing H_2_S donor sodium sulfide (Na_2_S), 1 µM-100 µM testosterone (T), and non-adrenergic, non-cholinergic nerve (NANC) mediated relaxation. NANC mediated relaxation was tested following a 30-minute incubation with 1 µM atropine and 30 µM Guanethidine followed by EFS. The following parameters were used for NANC EFS: frequencies of 2-128Hz (IIA and IPA) or 1-32Hz (CC), 10 second stimulation trains, 2 millisecond pulse width at 10 V (IIA and IPA) or 30 V (CC), and 2 minutes between each stimulation train. Relaxation responses were normalized as a percentage restoration to the resting tension from the PE pre-constricted value. Contractile responses were expressed as a percentage of maximal KCl-induced constriction. Pharmacologic dose-responses were fit with non-linear regression using GraphPad Prism. All vasoactive agents were dissolved in distilled water except for atropine, which was dissolved in 100% ethanol. Where atropine was added, the ethanol concentration in the bath was 0.1% (1:1000 dilution).

### 2.4 Immunoblotting Analysis

Separate cohorts of mice were used to obtain naïve CC tissue. Immediately following sacrifice, penile tissue was harvested from the base to the proximal glans, from which the corpus spongiosum, dorsal vein, and connective tissue were quickly stripped off. The corpus cavernosum was quickly rinsed in ice-cold PBS, blood was removed from the tissue, and the tissue snap frozen in liquid nitrogen and stored at -80°C until processing. Tissues were homogenized and immunoblotting analysis was performed as described [27]. Primary antibodies were obtained from Cell Signaling Technologies (CST; Danvers, MA, USA), or Protein Tech (PT; Rosemount, IL, USA) and used at the following dilutions: glutamate-cysteine ligase (Gclc, PT #12601-1-AP, 1:1000), optic atrophy type 1 (Opa1, CST #80471, 1:1000), peroxiredoxin 3 (Prdx3, PT #10664-1-AP, 1:1000), peroxiredoxin 5 (Prdx5, PT #17724-1-AP, 1:1000), mitofusin 1 (Mfn1, PT #13798-1-AP, 1:1000), mitofusin 2 (Mfn2, PT #12186-1-AP, 1:1000), NAD(P)H dehydrogenase quinone 1 (Nqo1, CST #62262, 1:1000), thioredoxin 1 (Trx1, CST #2298, 1:1000), thioredoxin-interacting protein (Txnip, PT #18243-1-AP, 1:1000), thioredoxin 2 (Trx2, CST #14907, 1:1000), dynamin-related protein 1 (Drp1, CST #8570, 1:1000), superoxide dismutase 1 (Sod1, PT #10269-1-AP, 1:1000), superoxide dismutase 2 (Sod2, CST #13141, 1:2000), superoxide dismutase 3 (Sod3, Santa Cruz Biotechnology #sc-271170, 1:1000) autophagy-related protein (Atg7, CST #8558, 1:1000), mitochondrial fission 1 (Fis1, PT #10956-1-AP, 1:1000), phosphorylated (Ser^403^) Sequestosome-1 (P-Sqstm1/p62, CST #39786, 1:1000), sequestosome-1 (Sqstm1/p62, CST #39749, 1:1000), heme oxygenase 1 (HO-1, PT #10701-1-AP, 1:1000), light chain 3B (LC3B, CST #2775, 1:1000), GAPDH (PT #60004-1-Ig, 1:4000). Anti-mouse (CST #7076, 1:4000) and anti-rabbit (Bethyl Laboratories #A120-102, 1:10000) secondary antibodies were used as appropriate.

### 2.5 Vascular Histology

Immediately following erectile function assessment IIA, IPA, and the penile shaft were dissected as previously described. Adherent fat and connective tissue were carefully excised under dissection microscope and once cleaned were placed upright in optimal cutting temperature (OCT, Sakura Finetek, Tokyo, Japan). Once in OCT the arteries and tissue were placed in 2-methylbutane supercooled in a dish of liquid nitrogen. Frozen OCT blocks were cut in 7 µM slices utilizing a microm HM 525 cryostat and placed on glass slides. Slides underwent Masson’s trichrome staining, which stains collagen blue and smooth muscle red, and were photographed with a digital microscope (LEICA DMi8) at a magnification of 20x (II and IPA) and 5 and 40x (CC). Images of tissue were analyzed for collagen to smooth muscle ratio via ImageJ software. Measurement of luminal size was achieved by mapping the inner wall of the artery and analyzing the mean area within the mapped wall. Measurement of smooth muscle was achieved by first mapping the external perimeter of the image then the lumen and finally the external edge of the artery is mapped excluding the blue stained sections. This mapped area is then deleted from the analysis highlighting only the remaining smooth muscle area. Finally, collagen is measured through the ImageJ software by adjusting the color threshold of the image to the full spectrum of blue and blue hues (values 140 to 175) and the resulting mean area is measured as collagen content. All histological analyses were performed in duplicate and the average values were reported.

### 2.6 Lipid assessment

Blood was obtained from mice in the freely fed state during the euthanasia process via cardiac puncture. Serum concentrations of total cholesterol, triglycerides, high-density lipoprotein cholesterol (HDL-C), low-density lipoprotein cholesterol (LDL-C), very low-density lipoprotein cholesterol (VLDL-C), alanine transaminase (ALT), and aspartate aminotransferase (AST) were obtained with a Piccolo Xpress chemistry analyzer (Abaxis, Union City, CA, USA) using lipid panel plus reagent disks (Abaxis #400-1030) for a subset of 10 mice/group. Serum samples were diluted 1:2 in phosphate buffered saline for analysis.

### 2.7 Statistical Analysis

Statistical analyses were performed with GraphPad Prism v10 (La Jolla, CA, USA). Statistical differences for the nerve-stimulated erectile response and the ex vivo vascular reactivity assessments were determined using a two-way repeated measures analysis of variance (ANOVA) followed by Tukey’s multiple comparisons post hoc analysis if main effects were detected. Group differences in metabolic characteristics, band intensity density for immunoblot images, and histologic staining differences were determined by one-way ANOVA followed by Tukey’s multiple comparisons post hoc analysis if main effects were detected. In cases where a Brown-Forsythe test determined significant differences in variances between groups, a Kruskal-Wallis test was used, followed by Dunn’s multiple comparisons post hoc analysis where significant differences were determined by the model. An α-level of 0.05 was used to determine statistical significance in all instances.

## Results

### 3.1 Metabolic Characteristics

Mice in the CLS group consumed an average of 18.8 ± 2.4 mg SG1002 per kg bodyweight per day (mg/kg/day), while mice in the CHS group consumed an average of 95.6 ± 8.7 mg/kg/day. Castration induced reductions in terminal body mass, total cholesterol, HDL, VLDL and triglycerides when compared to Sham (Table 1). Treatment with SG1002 did not significantly alter these decreases in lipid levels, although CLS exacerbated the castration-induced loss of body mass. CLS and CHS resulted in significant reductions in ALT that did not attain statistical significance in the Cast group when compared to Sham.

**Table 1:** Assessment of H_2_S supplementation on metabolic characteristics following castration. * p<0.05 vs Sham. ^π^ p<0.05 CLS vs Cast. Values are mean ± SD.

|  | Sham | Cast | CLS | CHS |
| --- | --- | --- | --- | --- |
| **Body Mass (g)** | 30.9 ± 3.0 | 28.4 ± 1.8* | 25.2 ± 2.0*<sup>π</sup> | 27 ± 2.6* |
| **Cholesterol (mg/dL)** | 105.3 ± 13.4 | 79.2 ± 8.9* | 75.4 ± 7.3* | 75.4 ± 7.1* |
| **HDL (mg/dL)** | 56.3 ± 8.2 | 41.8 ± 9.7* | 46.3 ± 3.6* | 45.3 ± 6.6* |
| **nHDLc (mg/dL)** | 48.4 ± 7.8 | 32.3 ± 8.2* | 28.8 ± 8.4* | 30 ± 5.7* |
| **LDL (mg/dL)** | 16.9 ± 10.3 | 12.2 ± 6.0 | 15.4 ± 6.1 | 10.7 ± 4.7 |
| **VLDL (mg/dL)** | 33 ± 7.35 | 21.6 ± 4.69* | 17.9 ± 7.77* | 19.3 ± 6.5* |
| **Triglycerides (mg/dL)** | 163 ± 38 | 107 ± 24* | 89 ± 39* | 97 ± 32* |
| **ALT (U/L)** | 63.3 ± 14.7 | 46.7 ± 20.4 | 33.6 ± 6.0* | 31.1 ± 8.9* |
| **AST (U/L)** | 87 ± 25 | 98 ± 35 | 93 ± 36 | 70 ± 11 |

### 3.2 SG1002 preserved erectile function in a castration model of androgen deprivation

Whether erectile function is assessed as peak ICP/MAP (Figure 1A) or as the AUC/MAP (Figure 1B), erectile function was diminished in all castrated cohorts at 1, 2 and 4 volts when compared to Sham. CLS mice saw preservation of erectile function for both the peak ICP/MAP and AUC/MAP measures in response to 1, 2, and 4 volts of electrical stimulation when compared to the Cast. However, erectile function was still impaired compared to the Sham controls. CHS mice saw preservation of erectile function for both the peak ICP/MAP and AUC/MAP measures throughout the voltage range of electrical stimulation when compared to Cast, although function remained significantly impaired relative to the Sham controls at the 1-4 volt stimulations.

**Figure 1:**
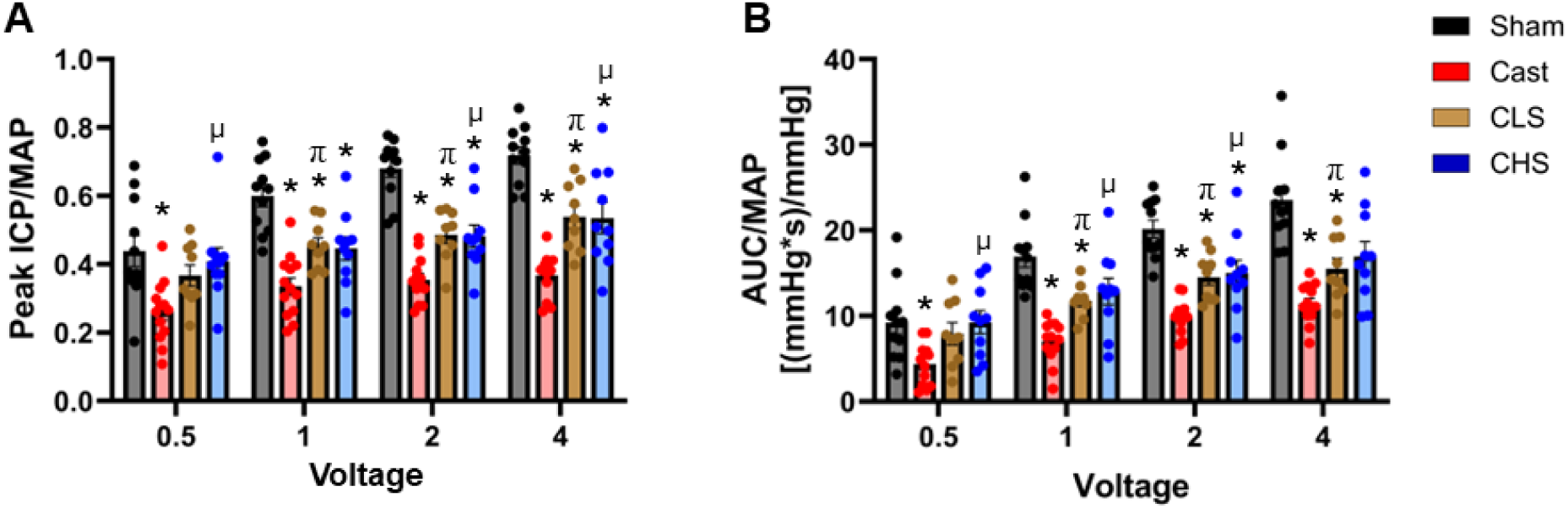
Assessment of erectile function via intracavernous pressure (ICP). Mice underwent sham surgery (Sham) or were surgically castrated (Cast). Two groups of castrated mice were treated with low-dose (CLS) or high-dose (CHS) SG1002. (**A)** Peak intracavernous pressure (ICP) normalized to mean arterial pressure (MAP) **(B)** Area-under-the-curve (AUC) normalized to MAP. Values represent means ± SEM for n = 9-13 animals per group. * p < 0.05 vs Sham, π p < 0.05 Cast vs CLS, µ p < 0.05 Cast vs CHS.

### 3.3 Effect of androgen deprivation and SG1002 treatment on ex vivo vascular relaxation

Endothelium-dependent relaxation as assessed via ACh application revealed significant effects of androgen deprivation on both IPA and CC relaxation at the four highest concentrations compared to the Sham controls (Figure 2A-C). Neither dose of SG1002 treatment afforded any protective effects against androgen deprivation. There were no significant effects of castration or SG1002 treatment observed in the IIA for this measure of endothelial function. Endothelium-independent relaxation as assessed by SNP application was significantly decreased in both the IPA and CC of all castrated groups throughout the majority of the effective concentration range relative to the Sham controls (Figure 2D-F). SNP-mediated relaxation was significantly impaired in the IIA of the Cast group at two SNP concentrations, while SG1002 treatment protected against this effect of androgen deprivation. Castration resulted in significant impairment in NANC-mediated relaxation in the CC as well as the IPA (Figure 2G-I). NANC-mediated relaxation of the CC was significantly reduced in all castrated groups throughout the 2-32 Hz frequency range. NANC-mediated relaxation of the IPA was significantly reduced in the Cast group at the 16, 64, and 128 Hz frequencies. While these responses were not significantly improved in the CHS group relative to Cast, they were not significantly impaired in the CHS group relative to Sham either.

**Figure 2:**
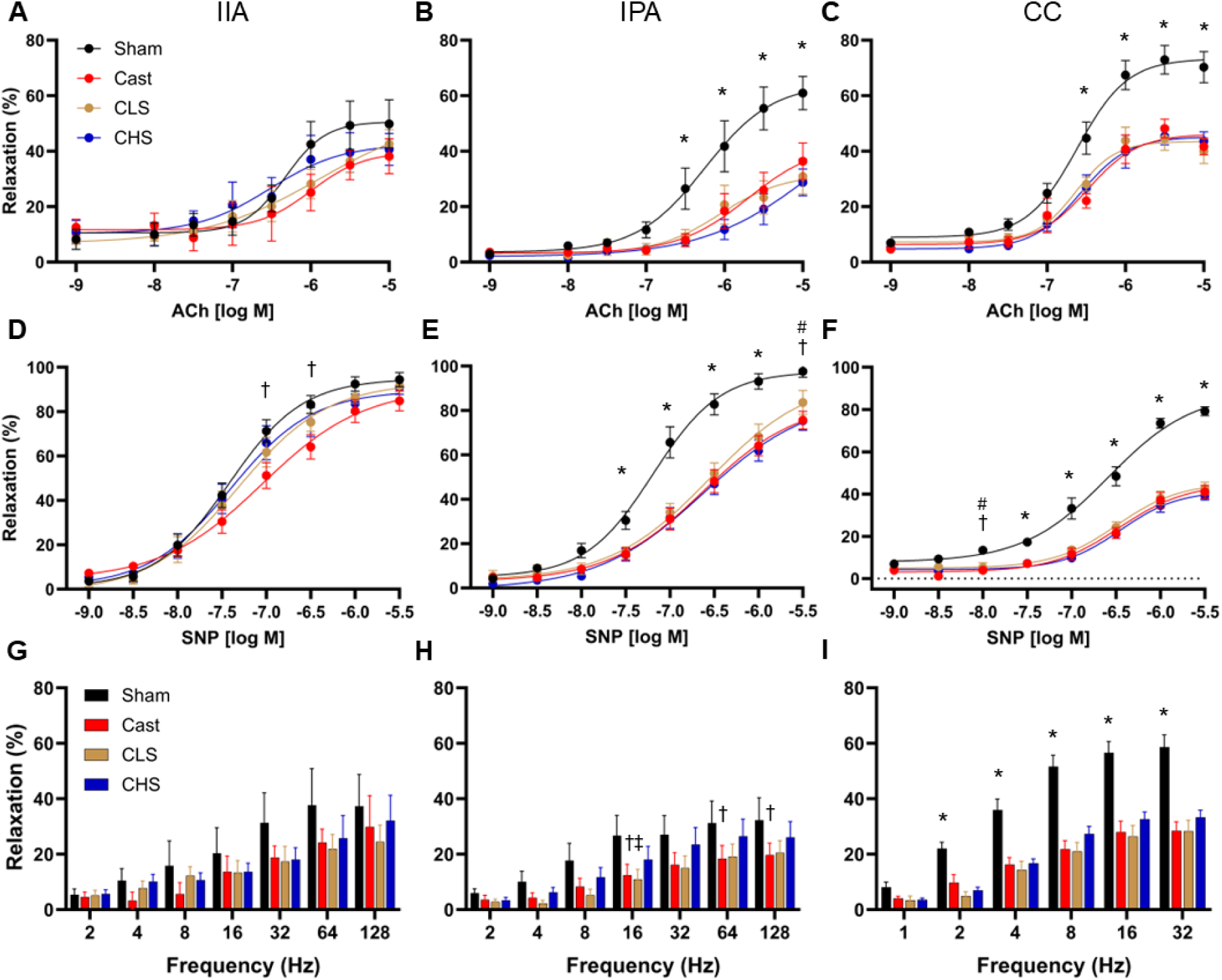
Assessment of endothelium-dependent, endothelium-independent, and NANC-mediated relaxation in the internal iliac artery (IIA), internal pudendal artery (IPA), and corpus cavernosum (CC) assessed via wire and muscle strip myography. Mice underwent sham surgery (Sham) or were surgically castrated (Cast). Two groups of castrated mice were treated with low-dose (CLS) or high-dose (CHS) SG1002. (**A-C**) Assessment of endothelial dependent relaxation induced via ACh (**D-F**) Assessment of endothelial independent relaxation induced via SNP. **(G-I)** Assessmnt of NANC-mediated relaxation. Values represent mean ± SEM for n = 5-13 animals per group * p < 0.05 Sham vs all castrated groups, † p < 0.05 Cast vs Sham, # p < 0.05 CHS vs Sham.

Testosterone induced vasodilation was significantly impaired in the CC and IPA of all androgen deprived groups, with the differences most evident in the middle of the dose-response curve (Figure 3A-C). There were no significant effects of SG1002 treatment on these responses. There were no significant alterations in testosterone-mediated relaxation in the IIA of any group. Vasodilation via H_2_S mediated hyperpolarization was assessed by application of the rapid releasing H_2_S donor Na_2_S in the organ bath. There were no significant differences observed amongst any group in the IIA (Figure 3D-F). The H_2_S-mediated vasorelaxation response of the IPA was similarly preserved in castrated mice, although there was a significant depression in the relaxation response at one point in the middle of the dose-response for castrated mice that were treated with the high-dose of SG1002. The H_2_S-mediated relaxation response of the CC was moderately but significantly augmented at one point in the middle of the dose-response curve in untreated castrated mice and castrated mice treated with the low-dose of SG1002 relative to the Sham controls.

**Figure 3:**
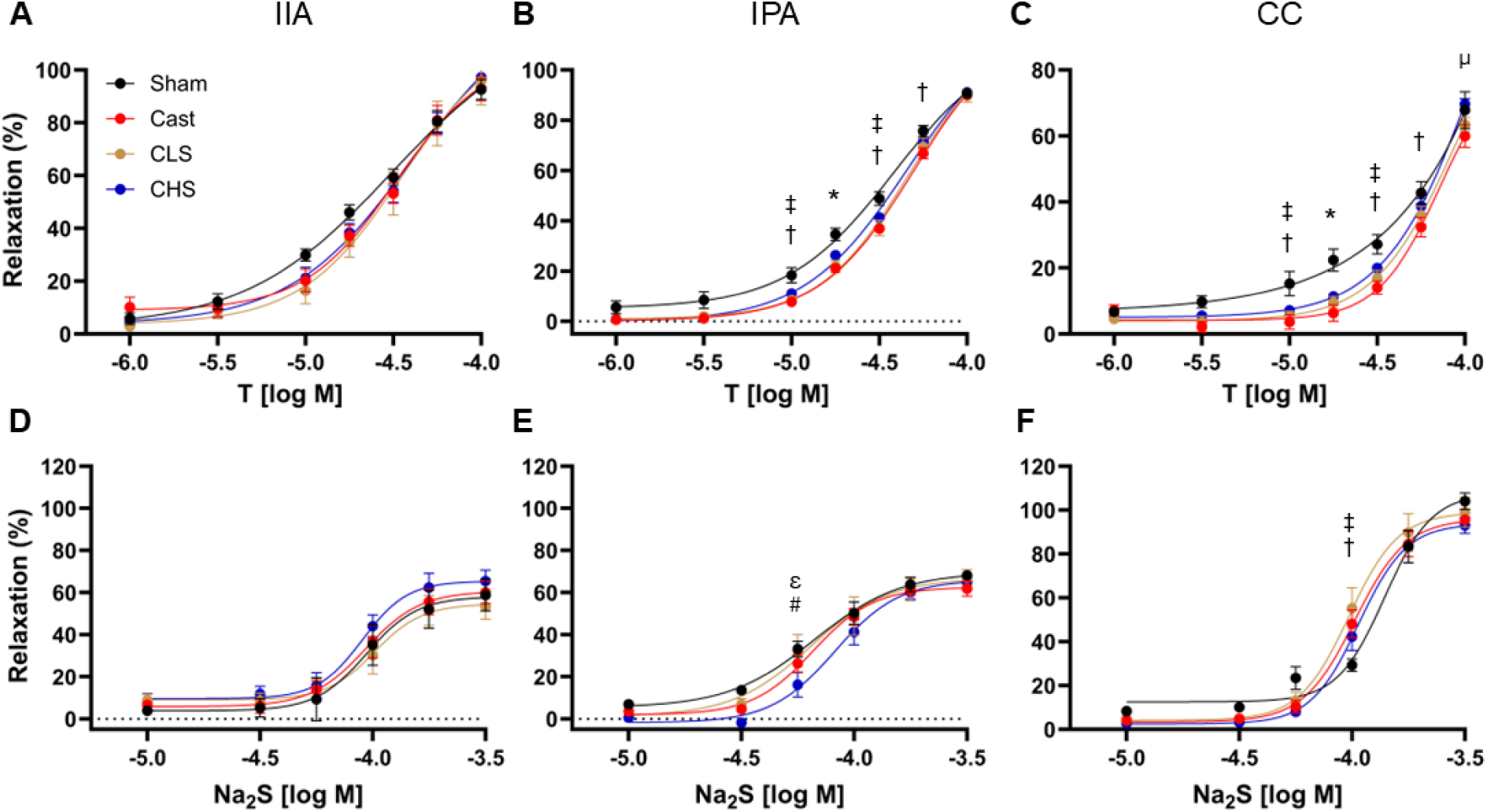
Assessment of testosterone-mediated and H_2_S-mediated vascular relaxation in the internal iliac artery (IIA), internal pudendal artery (IPA), and corpus cavernosum (CC) assessed via wire and muscle strip myography. Mice underwent sham surgery (Sham) or were surgically castrated (Cast). Two groups of castrated mice were treated with low-dose (CLS) or high-dose (CHS) SG1002. (**A-C**) Assessment of testosterone mediated relaxation induced via Testosterone. (**D-F**) Assessment of H_2_S mediated hyperpolarization induced via Na_2_S. Values represent mean ± SEM for n = 9-13 animals per group * p < 0.05 Sham vs all castrated groups, † p < 0.05 Cast vs Sham, ‡ p < 0.05 CLS vs Sham, µ p < 0.05 Cast vs CHS, ε p < 0.05 CLS vs CHS, # p < 0.05 CHS vs Sham.

### 3.4 Effect of androgen deprivation and SG1002 treatment on ex vivo vascular contraction

Androgen deprivation increased neurogenic vasoconstriction in all tissues investigated at the high end of the frequency range (Figure 4A-C). In the IIA, there were significant increases in constriction associated with castration at 16 and 32 Hz that were prevented by the low-dose of SG1002 treatment, but not the high-dose of treatment. Similarly, neurogenic vasoconstriction in the IPA was increased in the Cast group at 16 and 32 Hz relative to the Sham controls. This response remained significantly elevated in the CLS group at 32 Hz, while there was a trend toward elevation in the CHS group (p = 0.060) at 32 Hz. A significant increase in neurogenic constriction of the CC was observed for all castrated groups at 32 Hz, while the response at 16 Hz was also elevated at 16 Hz. Endothelin 1-mediated constriction of the IIA and IPA were not significantly altered by androgen deprivation or SG1002 treatment (Figure 4D-F). However, the contractile response to ET-1 in the CC was significantly elevated in the Cast group at one concentration and at four concentrations in the CLS group relative to the Sham controls.

**Figure 4.**
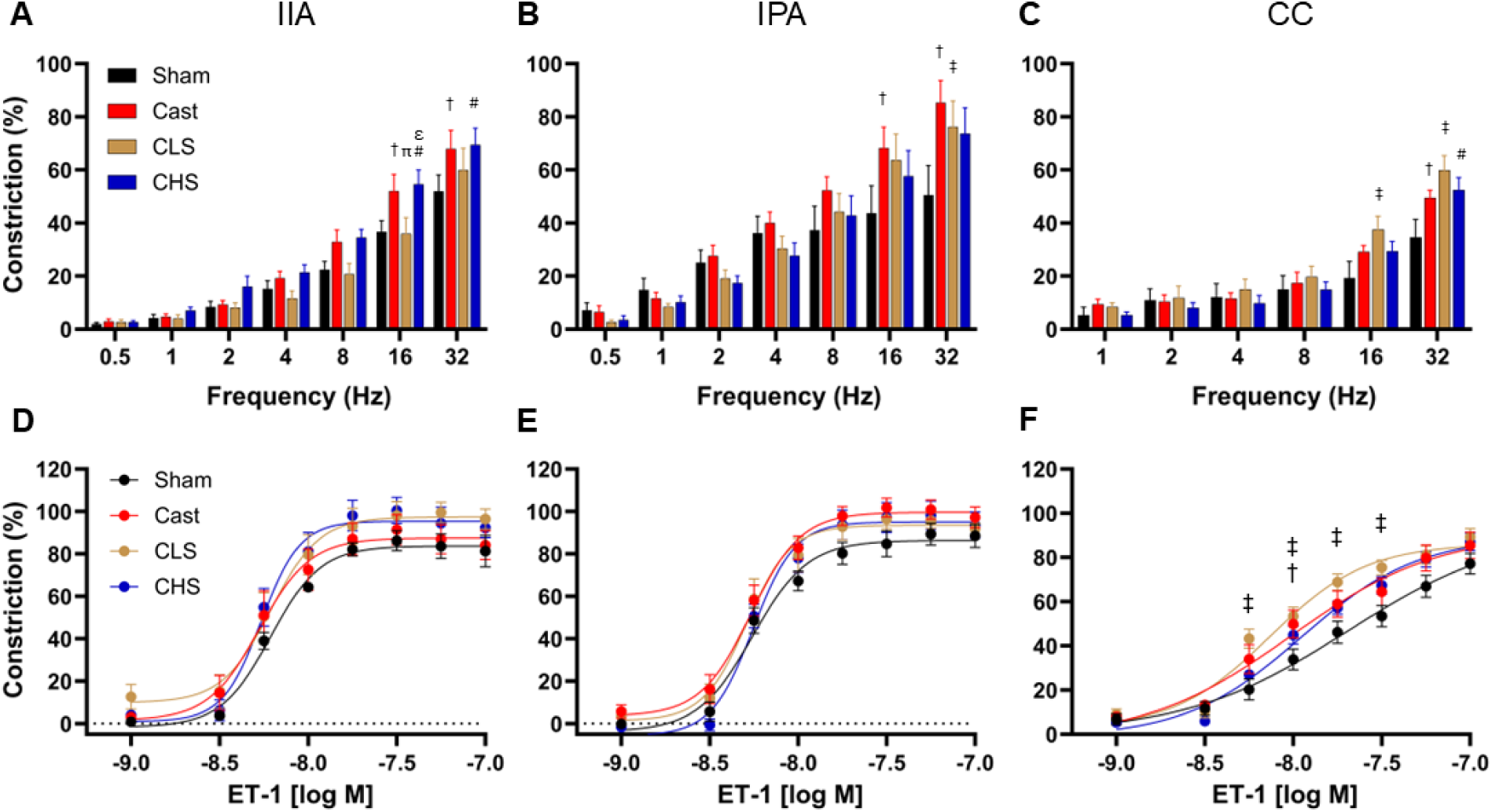
Assessment of neurogenic contraction and ET-1 mediated contractions in the internal iliac artery (IIA), internal pudendal artery (IPA), and corpus cavernosum (CC) assessed via wire and muscle strip myography. Mice underwent sham surgery (Sham) or were surgically castrated (Cast). Two groups of castrated mice were treated with low-dose (CLS) or high-dose (CHS) SG1002. (**A-C**) Assessment of electrical field stimulation-mediated contraction. (**D-F**) Assessment of ET1-mediated contraction. Values represent mean ± SEM for n = 9-13 animals per group. † p < 0.05 Cast vs Sham, ‡ p < 0.05 CLS vs Sham, # p < 0.05 CHS vs Sham, µ p < 0.05 Cast vs CHS, ε p < 0.05 CLS vs CHS, π p < 0.05 Cast vs CLS.

Androgen deprivation altered the α_1_-adrenergic vasocontractile profile across all tissues investigated (Figure 5A-C). The elevated IIA contractile response of the Cast group attained statistical significance relative to the Sham at the highest concentration of PE, although the responses of the CHS group were also elevated at two concentrations earlier in the dose-response curve. The PE-induced contractile responses in the IPA were significantly elevated in all castrated groups at the three highest concentrations with no effects of SG1002 treatment observed. PE-induced constriction of the CC was increased in the Cast group relative to the Sham group at two concentrations in the middle of the dose-response. However, this effect of androgen deprivation was completely suppressed by the high-dose SG1002 treatment, with the PE responses of the Cast group significantly higher than those of the CHS group at the five highest PE concentrations. Contractile responses to the thromboxane A_2_ receptor agonist U-46619 were also altered by androgen deprivation (Figure 5D-F). Contractile responses of the IIA were elevated in all castrated groups at two concentrations in the middle of the U-46619 dose-response curve. In the IPA, there was only one point in the dose-response where statistical significance was observed, with the CHS group demonstrating an elevated contractile response compared to the Sham controls. However, U-46619-mediated contraction of the CC was elevated at all but the lowest concentration of U-46619 tested in all castrated groups, with no effects of SG1002 treatment observed.

**Figure 5:**
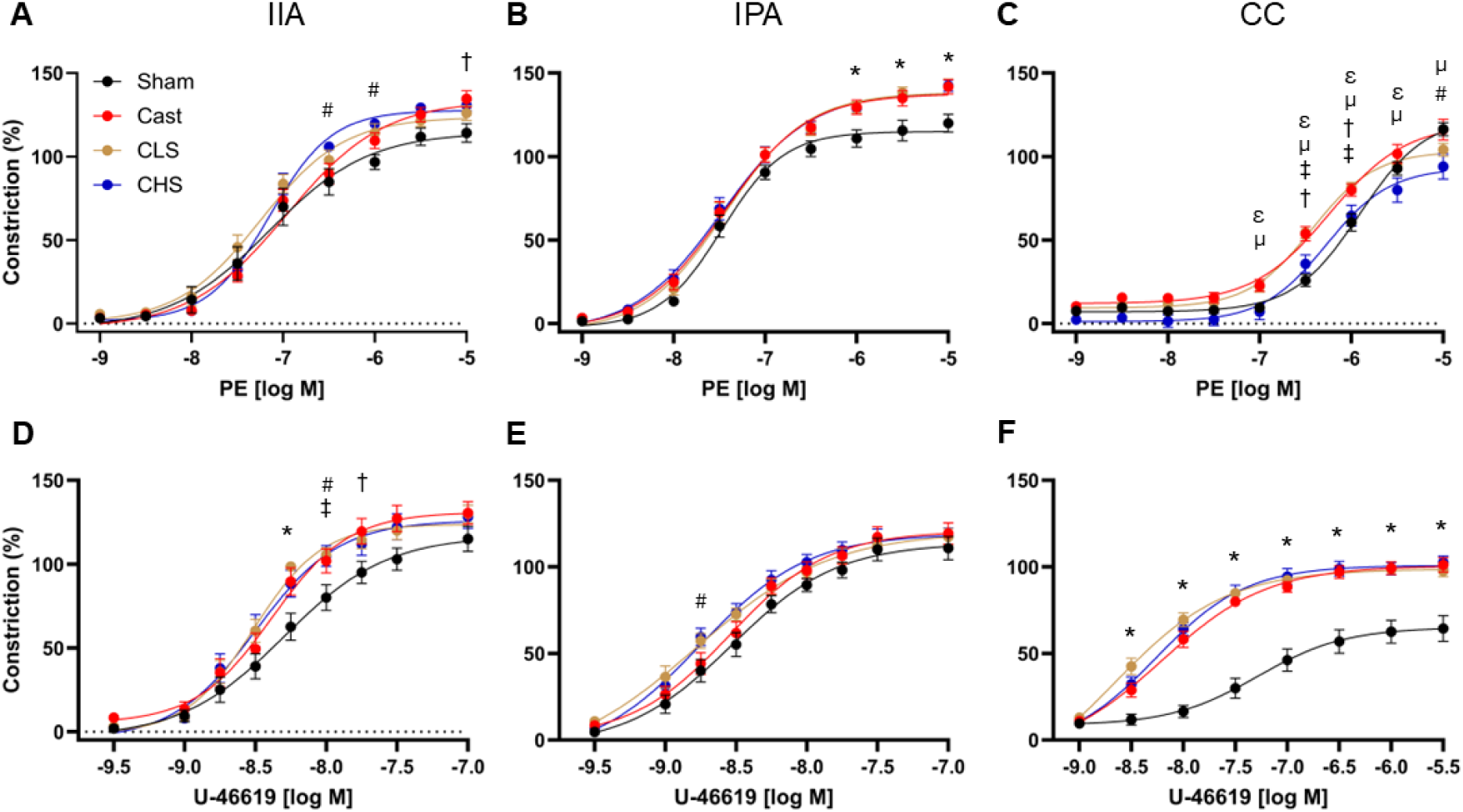
Assessment of α_1_-mediated and thromboxane A_2_ receptor-mediated contractions in the internal iliac artery (IIA), internal pudendal artery (IPA), and corpus cavernosum (CC) assessed via wire and muscle strip myography. Mice underwent sham surgery (Sham) or were surgically castrated (Cast). Two groups of castrated mice were treated with low-dose (CLS) or high-dose (CHS) SG1002. (**A-C)** Assessment of α_1_-mediated contraction induced via PE. (**D-F**) Assessment of thromboxane A_2_-mediated contraction induced via U-46619. Values represent mean ± SEM for n = 9-13 animals per group * p < 0.05 Sham vs all castrated groups, † p < 0.05 Cast vs Sham, ‡ p < 0.05 CLS vs Sham, # p < 0.05 CHS vs Sham, µ p < 0.05 Cast vs CHS, ε p < 0.05 CLS vs CHS.

### 3.5 Effect of androgen deprivation and SG1002 treatment on CC autophagic protein levels

Representative immunoblot images for LC3II, p62, P-p62, and Atg7 and their respective loading controls GAPDH are presented in Figure 6. Densitometry analysis (Figure 6) revealed no significant group differences for the autophagic markers of the LC3B ratio (p = 0.965), p62 (p = 0.129), or phosphorylated p62 (p = 0.512). There was however a downregulation of autophagic regulatory protein Atg7 (p < 0.001) with androgen deprivation, with post-hoc analysis confirming downregulation of Atg7 in Cast (p = 0.003), CLS (p = 0.007), and CHS (p < 0.001) relative to the Sham controls.

**Figure 6:**
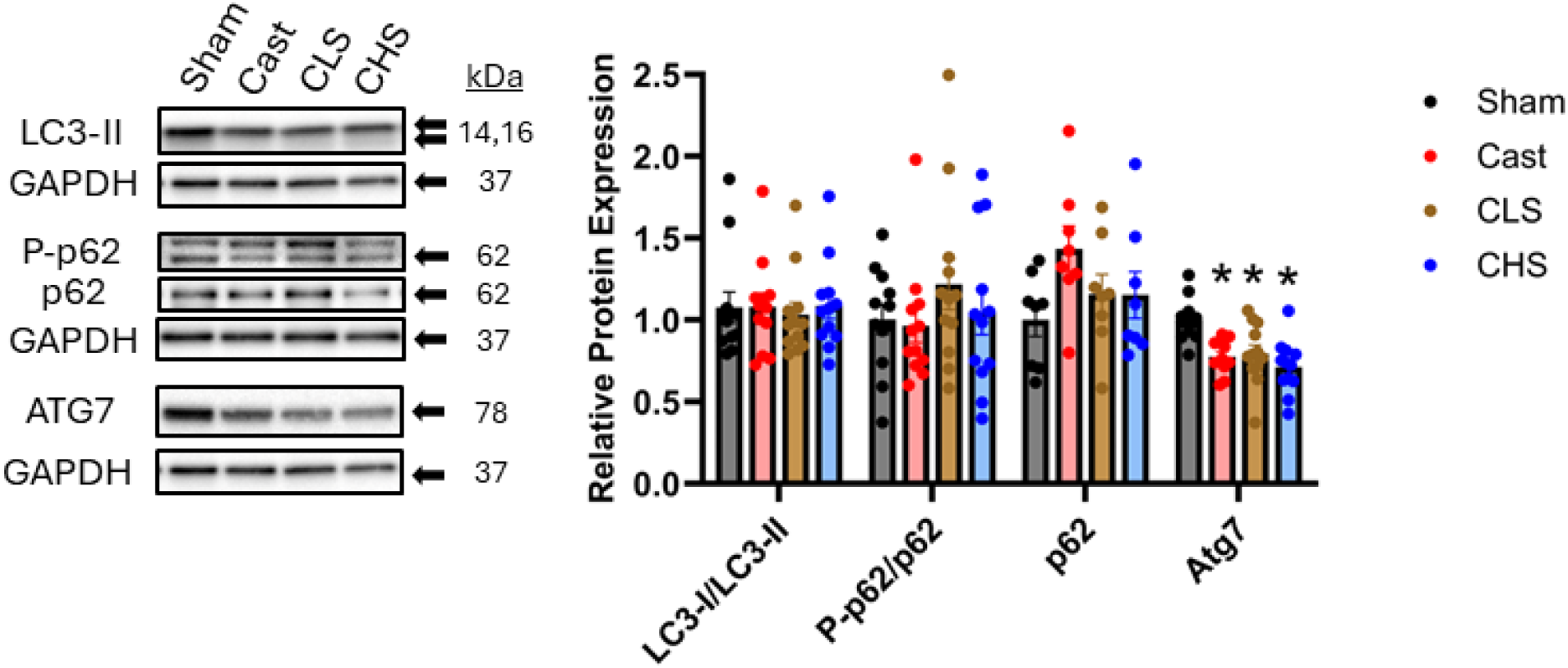
The expression of proteins related to autophagy in mouse corpus cavernosum measured by western blot. Mice underwent sham surgery (Sham) or were surgically castrated (Cast). Two groups of castrated mice were treated with low-dose (CLS) or high-dose (CHS) SG1002. Proteins include light chain 3B (LC3-II), P-p62, p62, and autophagy-related protein (ATG7). Protein expression was normalized to GAPDH or representative controls. Values represent means ± SEM for n = 12 animals per group. * p < 0.05 compared to Sham.

### 3.6 Effect of androgen deprivation and SG1002 treatment on CC levels of regulatory proteins of mitochondrial dynamics

Representative immunoblot images for Mfn1, Mfn2, Opa1, Drp1, Fis1 and their respective loading controls GAPDH are presented in Figure 7. Densitometry analysis (Figure 7) indicated a decreased expression of the mitochondrial fusion proteins Mfn1 (p = 0.014) and Mfn2 (p < 0.001) with androgen deprivation. Post-hoc analysis confirmed a decrease in Mfn2 in all castrated groups (Cast: p = 0.003, CLS: p = 0.005, CHS: p < 0.001) relative to Sham. Post-hoc analysis confirmed a decrease in Mfn1 of Cast (p = 0.023) and CLS (p = 0.046), although the decreased level of Mfn1 in the CHS group did not attain statistical significance (p = 0.157) compared to the Sham controls. There were no differences observed for the mitochondrial fusion protein Opa1 (p = 0.073), or the mitochondrial fission regulatory proteins Drp1 (p = 0.456) or Fis1 (p = 0.907).

**Figure 7:**
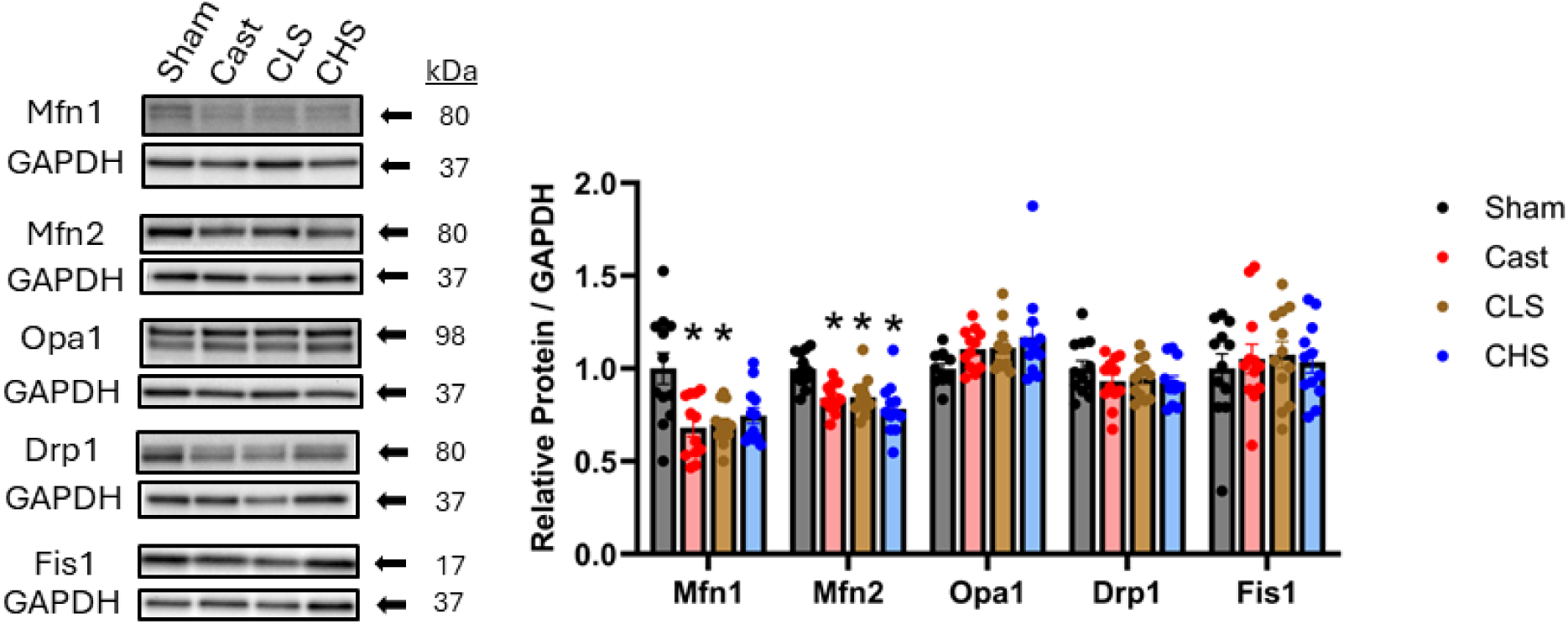
The expression of proteins related to mitochondrial dynamics in mouse corpus cavernosum measured by western blot. Mice underwent sham surgery (Sham) or were surgically castrated (Cast). Two groups of castrated mice were treated with low-dose (CLS) or high-dose (CHS) SG1002. Proteins include mitofusin 1 (Mfn1), mitofusin 2 (Mfn2), optic atrophy type 1 (Opa1), dynamin-related protein 1 (Drp1), and mitochondrial fission 1 (Fis1). Protein expression was normalized to GAPDH. Values represent means ± SEM for n = 12 animals per group. * p < 0.05 compared to Sham.

### 3.7 Effect of androgen deprivation and SG1002 treatment on CC cellular antioxidant defense proteins

Representatives immunoblot images for, Trx1, Trx2, Txnip, Prdx3, Prdx5, and their respective loading controls GAPDH are presented in Figure 8. Densitometry analysis (Figure 8) revealed decreased expression of the cytosolic redox regulatory protein Trx1 (p < 0.001) with androgen deprivation. Post-hoc analysis confirmed differences in Cast (p = 0.029), CLS (p = 0.003), and CHS (p = 0.002) compared to the Sham control. There were no differences observed for the mitochondrial redox regulatory protein Trx2 (p = 0.985), or the mitochondrial antioxidant enzymes Prdx3 (p = 0.167) or Prdx5 (p = 0.164). There was however a decrease in Txnip levels that was only present the CHS group compared to the Sham group (p = 0.033). Txnip helps regulate cellular oxidant stress through inhibiting activity of the thioredoxin redox systems, thus a downregulation of Txnip associated with the high-dose of SG1002 treatment may represent a mechanism through which H_2_S therapy could suppress oxidative stress.

**Figure 8:**
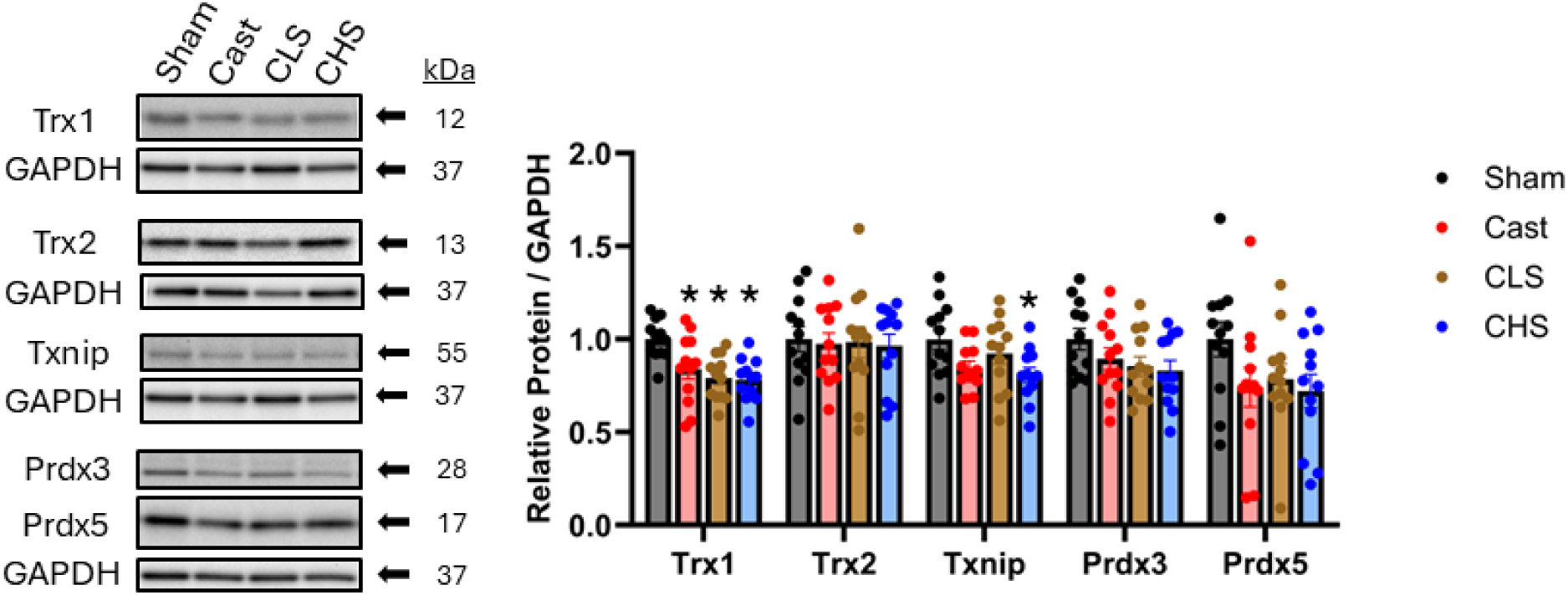
The expression of proteins related to the thioredoxin system in mouse corpus cavernosum measured by western blot. Mice underwent sham surgery (Sham) or were surgically castrated (Cast). Two groups of castrated mice were treated with low-dose (CLS) or high-dose (CHS) SG1002. Proteins include thioredoxin 1 (Trx1), thioredoxin 2 (Trx2), thioredoxin-interacting protein (Txnip), peroxiredoxin 3 (Prdx3), peroxiredoxin 5 (Prdx5). Protein expression was normalized to GAPDH. Values represent means ± SEM for n = 12 animals per group. * p < 0.05 compared to Sham.

Representative immunoblot images for Gclc, Nqo1, Sod1, Sod2, Sod3, HO-1, and their respective loading controls GAPDH are presented in Figure 9. Densitometry analysis (Figure 9) revealed decreased expression of Gclc with androgen deprivation (p < 0.001), which is a key enzyme in the biosynthesis of glutathione. Post-hoc analysis confirmed decreases in Cast (p = 0.007), CLS (p < 0.001), and CHS (p = 0.015) compared to the Sham controls. There were no significant differences observed for Nqo1 (p = 0.985), the mitochondrial dismutase Sod2 (p = 0.830), or HO-1 (p = 0.077). Interestingly, there was an increased expression of the cytosolic dismutase Sod1 in the Cast (p = 0.002) and CHS (p = 0.034) groups relative to the Sham control. However, there was a major decrease in the extracellular dismutase Sod3 of all androgen deprived groups (p < 0.0001) compared to the Sham control. The mean levels of Sod3 in the CHS group were higher than the Cast group, though this did not reach statistical significance (p = 0.132) following post-hoc analysis.

**Figure 9:**
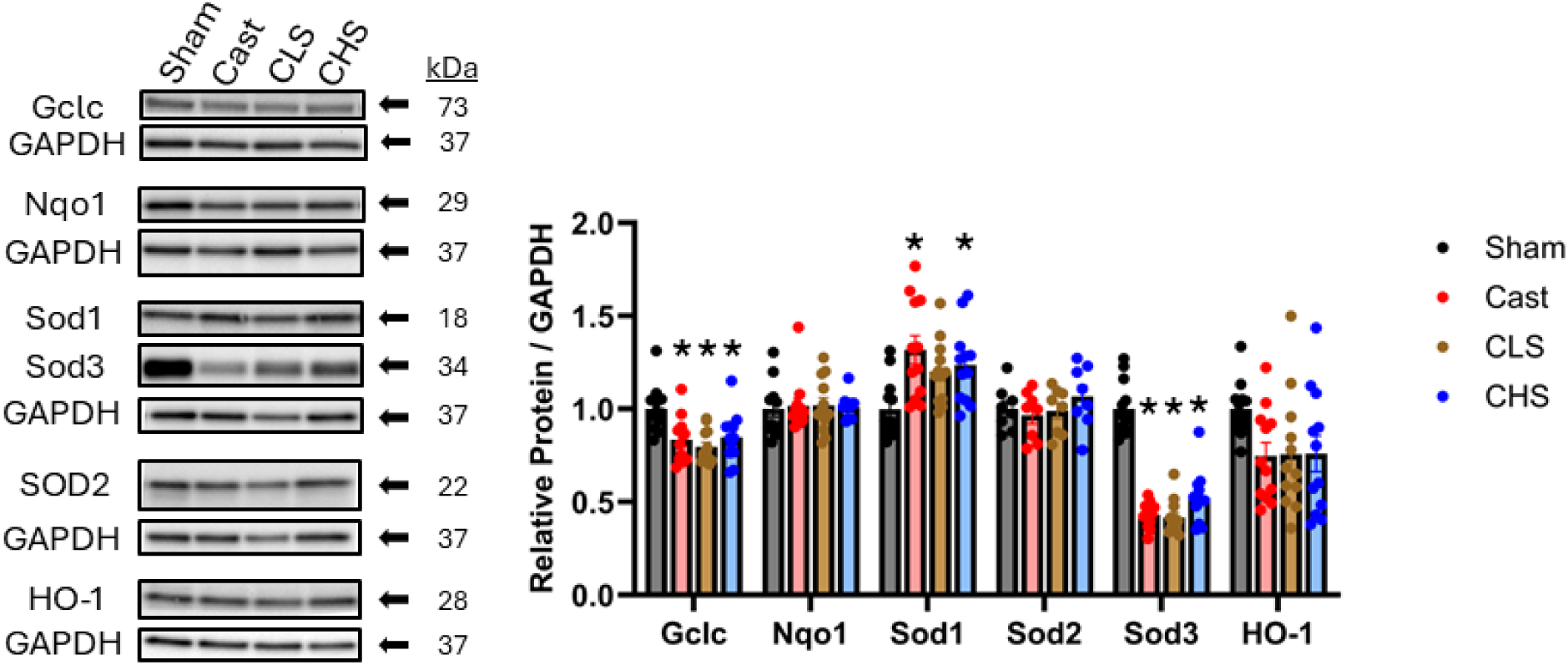
The expression of proteins related to oxidant dismutation and detoxification in mouse corpus cavernosum measured by western blot. Mice underwent sham surgery (Sham) or were surgically castrated (Cast). Two groups of castrated mice were treated with low-dose (CLS) or high-dose (CHS) SG1002. Proteins include glutamate-cysteine ligase (Gclc), NAD(P)H dehydrogenase quinone 1 (Nqo1), superoxide dismutase 1 (Sod1), superoxide dismutase 2 (Sod2), superoxide dismutase 3 (Sod3), heme oxygenase 1 (HO-1). Protein expression was normalized to GAPDH. Values represent means ± SEM for n = 12 animals per group. * p < 0.05 compared to Sham.

### 3.8 Androgen deprivation induces fibrotic remodeling of the IPA and CC

Representative images for Masson’s trichrome staining of the IIA, IPA, and CC are presented in Figure 10 (A-D). Quantifications of the smooth muscle:collagen ratio are depicted in Figure 10E. There were no differences amongst any group for the IIA (p = 0.856). There was a decrease in smooth muscle:collagen in the IPA with androgen deprivation (p = 0.026), with post-hoc analysis revealing a significant decrease only in the Cast group relative to the Sham group (p = 0.017). Similarly, there was a decrease in smooth muscle:collagen in the CC with androgen deprivation, with post-hoc analysis revealing a significant decrease only in the Cast group relative to the Sham group (p = 0.024), although there was a trend observed for the CLS group (p = 0.065) compared to the Sham group.

**Figure 10:**
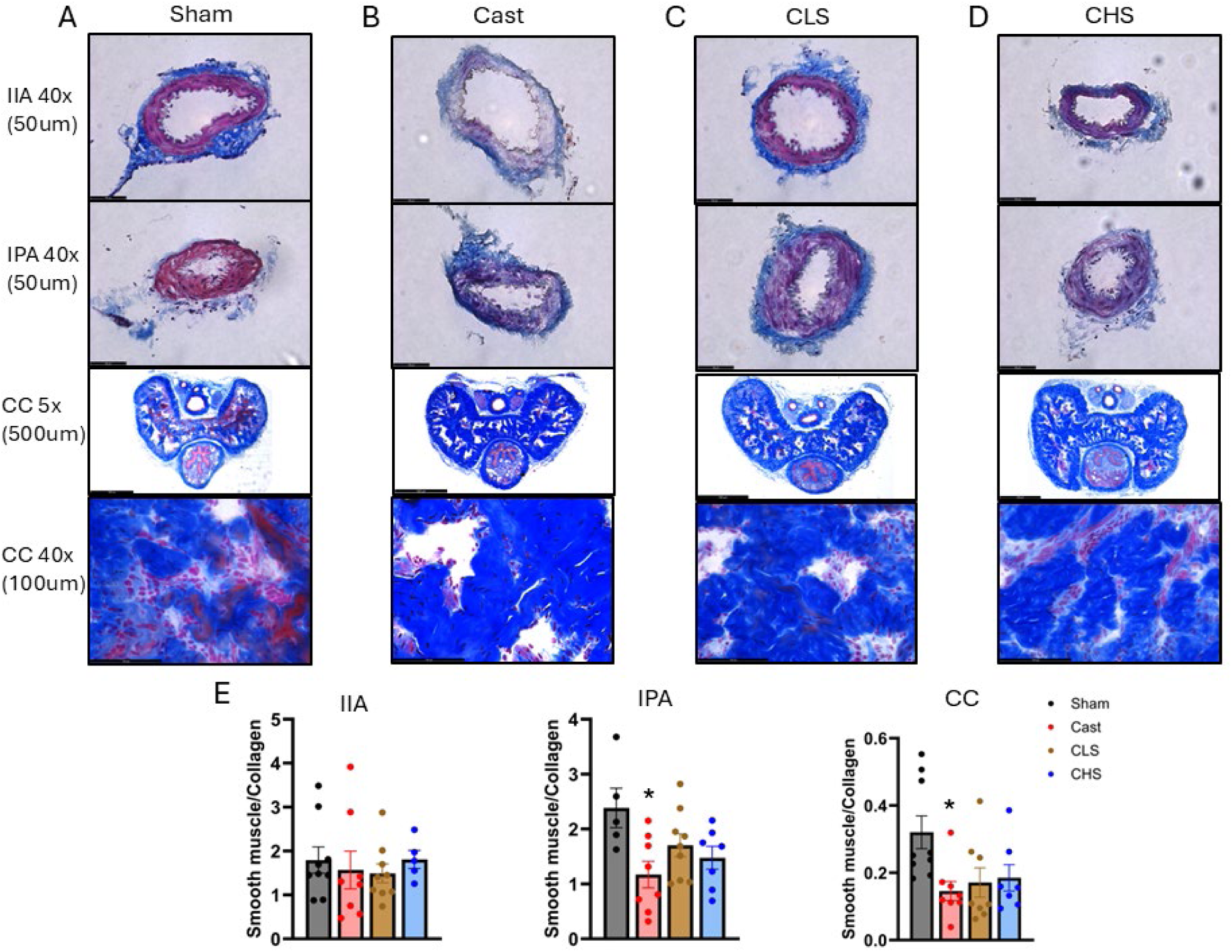
Assessment of area of smooth muscle compared to collagen area in the internal iliac artery (IIA), internal pudendal artery (IPA), and corpus cavernosum (CC). **(A)** Mice underwent sham surgery (Sham) or **(B)** were surgically castrated (Cast). **(C-D)** Two groups of castrated mice were treated with low-dose (CLS) or high-dose (CHS) SG1002. **(E)** Smooth muscle to collagen ratios were assessed utilizing ImageJ software. Images of both IIA and IPA were taken at 40x magnification. Images of penile cross section were taken at 5x magnification. Close up images of the corpus cavernosum were taken at 40x magnification. Values represent means ± SEM for n = 5-9 animals per group * p<0.05 vs Sham.

## 4. Discussion

ADT has continually shown to alter sexual functioning and increase the risk and rate of ED [3,28]. The present study found markedly impaired erectile function in castrated mice across the voltage range regardless of whether erectile function is assessed by peak ICP/MAP or AUC/MAP (Figure 1), indicating severe impairments in both the ability to achieve and maintain an erection. This decrease in erectile function as measured via ICP/MAP has been similarly seen by other laboratories in castrated rat models [29–31]. The two doses of SG1002 treatment exerted similar effects, whereby erectile function was improved relative to the untreated castrated group, but still significantly impaired relative to the sham group. Looking at the two highest stimulation parameters (2 and 4V) between both ICP/MAP and AUC/MAP measures, these improvements translate to 34-49% and 39-50% mean preservations of function from the castrated values to the sham values for CLS and CHS, respectively. As the degree of ED following ADT is often severe, such improvements may not be sufficient to fully preserve satisfactory sexual function to the patient. However, it is important to note that current first line therapies (i.e. PDE5 inhibitors) also have poor efficacy in this patient population [32]. It is plausible such improvements exerted by H_2_S therapy could improve the efficacy of PDE5 inhibitors, although further research will be needed to address the potential of this combinatorial approach.

We performed myography on the IIA, IPA, and CC to gain further insight into the mechanisms responsible for these functional alterations, as well as the relative contribution of each of these tissues involved in the erectile process. We observed marked impairments in endothelial function of the IPA and CC following castration, as assessed by ACh-mediated relaxation (Fig. 2A-C). However, we also observed a stark decrement in the relaxation response to the NO-donor SNP (Fig. 2D-F), thus the true impairment is more likely in the ability of the smooth muscle to respond to NO rather than the ability of the endothelium to produce NO. Neither dosing paradigm of SG1002 had any effect on these outcomes in the IPA or CC. However, we observed a more modest decrement in SNP-mediated relaxation of the IIA that was mostly prevented by SG1002 treatment, indicating that this could be contributing factor to the protective effects of SG1002 on erectile function. Moreover, NANC nerve-mediated relaxation was robustly impaired in the CC following castration and more modestly, but significantly, impaired in the IPA (Fig. 2G-I). NANC nerve activation and the subsequent release of NO play a key role in the mechanism of smooth muscle relaxation and subsequent erectile function [33].

However, given the drastically impaired responsiveness of the smooth muscle to the NO-donor in these tissues following castration, we again cannot be certain that there is an impairment in NANC nerve transmission. Interestingly, there was a moderate preservation of NANC nerve-mediated relaxation in the IPA of castrated mice treated with the high dose of SG1002, which could potentially reflect a small effect of H_2_S therapy on neurotransmission in these arteries.

Alves-Lopes et al. previously observed a reduction in NANC-mediated relaxation of the IPA of rats in response to a slightly shorter duration of androgen deprivation [6]. They observed a preserved relaxation response to SNP in the IPA, making a more clear demonstration that NANC neurotransmission is impaired in these arteries following castration [6].

The vasodilatory response to testosterone was also significantly impaired in the IPA and CC following castration (Fig. 3A-C), albeit to a much lesser degree than the NO-reliant responses. This is particularly of interest as mice have shown to have a reflexive testosterone release upon sexual arousal, meaning the immediate effect of a pulsatory increase in testosterone has direct implications for sexual function [34,35]. Although the depressive effects of ADT on libido are well-documented, loss of the vasodilatory effects of testosterone may be another minor contributing factor to ED in these patients, while impaired sensitivity to the vasodilatory effects of testosterone in these tissues may be yet another factor to contend with upon re-androgenization of the tissues following cessation of ADT. An interesting finding of this study was that while all NO-mediated mechanisms of relaxing the castrated IPA and CC were markedly impaired, there were no impairments in the vasodilatory response to the rapid releasing H_2_S donor Na_2_S in these tissues (Fig. 3D-F). Remarkably, there was an augmented relaxation in the castrated CC in the middle of the Na_2_S dose-response curve. We have previously observed the phenomenon in healthy tissues where this H_2_S donor exhibits greater relaxation potential in the CC than the IIA or IPA, which is the opposite of the relaxation profile of the NO donor SNP [36]. That H_2_S has the greatest relaxant effects in the tissue most affected by androgen deprivation (CC) has important implications. First, the *in vivo* impact of SG1002 therapy on partially preserving erectile function following castration could be due to local actions of H_2_S in the CC acting to reduce resistance to blood flow into the end organ. Second, harnessing the mechanism of H_2_S action for localized delivery, such as an intracavernous injection, could conceivably improve efficacy of injectable therapies in ADT patients.

Increases in contractility of the CC and/or the pre-penile arteries leads to a reduction in smooth muscle relaxation that is critical to healthy erectile functioning. The penis is maintained in the flaccid state by various mechanisms of vasoconstriction, of which the vasodilatory mechanisms must overcome to induce an erection. We observed increases in neurogenic constriction following castration at the higher end of the frequency range for all tissues (Fig. 4A-C), with no consistent effects of SG1002 therapy. ET-1 release from endothelial cells is another contributing factor to maintaining the penis in a flaccid state. While we cannot verify whether or not local ET-1 release was altered by castration or H_2_S therapy, there was a slight increase in the contractile response to ET-1 observed in the CC following castration (Fig. 4D-F). This effect was slightly exacerbated by the low-dose of SG1002 treatment. Perhaps the most significant contributor to maintaining the flaccid penis is the sympathetic nervous system, for which the vasoconstrictive actions are predominantly carried out by activation of α_1_-adrenoceptors [37].

Our dose-response applications of the α_1_-agonist PE revealed heightened vasoconstriction in the IIA, IPA, and CC following castration (Fig. 5A-C), with the effects in the CC and IPA being apparent through a wider range of the dose-response curve than the IIA. Treatment with the high dose of SG1002 significantly suppressed PE-mediated contraction relative to the untreated castrated group in the CC at the five highest doses of PE tested. This decrease in responsiveness to α_1_-receptor activation in the CC induced by H_2_S therapy could be another contributing factor to the observed partial preservation of *in vivo* erectile function. Our previous study revealed heightened PE-constriction in the CC 8-weeks following castration in the middle of the PE response curve, similar to the present study. However, we previously observed no differences in PE-constriction in the IPA following castration [5]. These findings in the IPA were thus unexpected, and could potentially be a function of the shortened period of androgen deprivation in the present study.

We observed a significant increase in constriction of the IIA of all castrated groups in response to the thromboxane A_2_ (TXA_2_) receptor agonist U-46619, but perhaps the most stunning findings of the present study are the magnitude of elevated contractions of the androgen deprived CC in response to this agonist (Fig. 5D-F). TXA_2_ is most notably produced by aggregated platelets, where it has a very short half-life and acts locally near its site of production [38]. Although these *ex vivo* results are not indicative of *in vivo* TXA_2_ production, they clearly indicate that TXA_2_ produced in the CC will have a greater impact on impeding erections in the androgen deprived state. NO and prostacyclin notoriously prevent platelet aggregation and subsequent TXA_2_ production, but stimuli that promote platelet derived TXA_2_ production include inflammation, vascular damage and endothelial cell injury, elevated ROS levels, and exposure of platelets to collagen [39]. The androgen deprived CC may provide an environment suitable for rampant TXA_2_ production. Moreover, androgen deprived platelets themselves may be more prone to aggregation. Circulating concentrations of a stable TXA_2_ metabolite have been shown to be elevated in prostate cancer patients undergoing ADT, as well as in rats following castration [40,41]. Furthermore, exposure of platelets to physiological androgen levels in an *in vitro* setting suppresses ADP- and H_2_O_2_-stimulated platelet aggregation compared to androgen deprived platelets [41]. While more work is needed to ascertain the potential influence of H_2_S therapy on platelet aggregation pertinent to erectile dysfunction, *in vitro* exposure of platelets to H_2_S prevents aggregation in response to a variety of aggregatory stimuli in a dose-dependent manner [42,43].

Autophagy is the self-degradative process by which homeostasis is maintained at a cellular level by the breakdown and removal of damaged proteins and organelles and recycling of intracellular components [44]. Mitophagy is the specific autophagic process that takes place within the mitochondria [45]. While autophagy and mitophagy have been demonstrated to have a role in maintaining vascular and smooth muscle health and function their causal role in erectile dysfunction is still being elucidated. We previously demonstrated the protective effect of the HDAC6 inhibitor, tubacin, on vascular and erectile health independent of changes in mitophagy and autophagy in a hypercholesterolemic mouse model [46]. However, Zheng et al. found that improvement in erectile function seen in type 2 diabetic rats given Icariside II was potentially via the blunting of the excessive mitochondrial autophagy in the SMCs of the corpus cavernosum [47]. In the present study, castration did not result in significant reductions in protein expression for autophagy as assessed via LC3-II and P-p62 (Fig. 6). However, castration did result in a significant reduction in expression of Atg7 (Fig. 6). Atg7 plays an essential role in regulating both LC3-II and Atg12-Atg5 autophagic pathways. The impairment or deletion of Atg7 causes a significant disruption in autophagy and has been utilized to demonstrate the role of autophagy in the manifestation and progression of various disease states, including the growth of prostate cancer tumors [48–53].

Androgen deprivation also altered CC levels of proteins that regulate mitochondrial dynamics, specifically through a decrease in the mitochondrial fusion regulatory proteins Mfn1 and Mfn2 (Fig. 7). The high-dose of SG1002 treatment partially prevented the suppressive effects of androgen deprivation on Mfn1, although it had no effect on Mfn2 levels. Suppression of either Mfn1 or Mfn2 results in a negative cascade of mitochondrial dysfunction via deregulating mitochondrial fusion and impairment of cellular proliferation and ROS clearance [54,55]. Impairments in mitochondrial fusion have also been attributed to phenotypic switches in smooth muscle resulting in changes from the contractile to synthetic state [56]. While H_2_S has been shown to play a role in the regulation and maintenance of mitophagy and mitochondrial dynamics in other tissues such as cardiac muscle of db/db mice [15], the effects of SG1002 treatment on proteins related to autophagy or mitochondrial dynamics in the androgen deprived CC were marginal to negligible.

Li et al. have previously demonstrated that castration drastically increased ROS production in the CC of castrated rats. Additionally, administration of exogenous testosterone was able to reduce this spike in ROS production [29]. Increased ROS will result in oxidative stress and subsequent cellular damage and dysregulation of cell signaling [57]. H_2_S acts in an antioxidative capacity via its ability to scavenge ROS and its role in Nrf2 signaling through its ability to sulfhydrate Keap1 [58]. Kaep1 binding to Nrf2 maintains Nrf2 in an inactive complex. Upon sulfhydration of Keap1, Nrf2 is released from Kaep1 allowing for the binding of Nrf2 to promotors of antioxidant response elements (AREs) [10,59]. Androgen deprivation resulted in detrimental alterations in antioxidative capacity via downregulation of Gclc, Trx1, and Sod3 (Fig 8-9), with no impact of SG1002 treatment observed. We have previously found that the low-dose SG1002 treatment stimulates CC expression of Nqo1, Gclc, Prdx3, and HO-1 in intact mice [24]. It appears that androgenization of the tissues is necessary for these stimulatory effects to occur. While there were no substantial effects of SG1002 treatment on expression of proteins involved in ROS quenching and cellular antioxidant defense, we cannot be certain that SG1002 did not impact penile ROS balance through direct H_2_S driven ROS scavenging.

Androgen deprivation induced decreases in total cholesterol and triglycerides, as well as HDL, nHDLc, and VLDL (Table 1). These alterations in lipid profiles differ from those commonly seen in individuals undergoing ADT with most studies finding increases in triglyceride and cholesterol levels [60]. SG1002 treatment had no apparent adverse effects on lipid profiles at either dose. As expected, androgen deprivation decreased bodyweight in all castrated groups when compared with Sham controls (Table 1), likely through decrease in muscle mass. Moreover, androgen deprivation induced a significant loss in the smooth muscle to collagen ratio in the IPA and CC (Fig 10). SG1002 treatment provided a modest protective effect against fibrotic remodeling in the IPA, with mean values representing 44% and 25% preservations of the smooth muscle to collagen ratio for the low-dose, and high-dose treatments, respectively. Decreases in endogenous H_2_S have been found to promote a pro-fibrotic environment making H_2_S therapies a target for combating these compositional changes [61,62]. While adverse tissue remodeling remained evident despite SG1002 treatment in these tissues that are critical for satisfactory erectile function, it is important to note that ADT is not permanent, thus any preservation of tissue function or quality could result in better patient outcomes upon cessation of ADT and reandrogenization of these tissues.

There were several limitations to the present study. Although we designed the low-dose of SG1002 treatment to be identical to prior studies that have demonstrated a clear increase in circulating free sulfide and sulfane sulfur levels with SG1002 administration [22,23], we were unable to perform such measures for this study. While we did not observe any clearly adverse effects of this or the 5x higher dose of SG1002 treatment on erectile or metabolic health, inclusion of a control group treated with these doses of SG1002 would further establish the safety profile of these treatment regimens. We were further unable to measure penile ROS levels. While our data firmly allows us to reject our hypothesis that SG1002 would stimulate CC expression of antioxidant proteins, we cannot be certain that SG1002 did not affect the penile redox environment through direct oxidant scavenging of H_2_S. Finally, due to technical errors in processing the tissues for histology, our sample sizes for some of the histological measures were smaller than desired. SG1002 treatment provided several marginally protective effects that collectively resulted in a modest, but potentially meaningful retention of erectile function following the long-term androgen deprived period. From a clinical translation standpoint, such protection could result in a higher efficacy of PDE5 inhibitors during the ADT period, or a faster restoration of erectile function following cessation of ADT, either with or without the use of PDE5 inhibitors. Future research could test these possibilities in two ways. First, a combinatorial treatment strategy could be tested with administration of SG1002 and a PDE5 inhibitor throughout the androgen deprived period. The potential of added blood perfusion of the CC induced by the PDE5 inhibitor could preserve smooth muscle structure and function. Second, this SG1002 treatment paradigm as well as the combinatorial strategy could be tested with a chemical castration approach that is more similar to ADT, with mice tested at various timepoints following cessation of the ADT period.

## 5. Conclusion

SG1002 treatment provided a moderate but potentially clinically meaningful preservation of erectile function following a prolonged state of androgen deprivation. Androgen deprivation induced major functional alterations including impaired vasorelaxation and increased vasoconstriction in the erectile tissue of the corpus cavernosum and internal pudendal artery, whereas the effects in the internal iliac artery were more modest. SG1002 treatment was largely ineffective at preventing androgen deprivation-induced alterations in proteins related to cellular autophagy and antioxidant defense in the corpus cavernosum, as well as fibrotic remodeling of the corpus cavernosum, but partially prevented fibrotic remodeling of the internal pudendal artery.

## CRediT authorship contribution statement

CMI: methodology, validation, formal analysis, investigation, visualization, data curation, writing – original draft.

CJP: methodology, validation, formal analysis, investigation, data curation, project administration.

TAA: methodology, validation, investigation.

JDL: conceptualization, investigation, project administration, funding acquisition, supervision, writing – review and editing.

## Funding

JDL is supported by grant K01DK115540 from the National Institutes of Health. This study was funded by grant R03DK131242 from the National Institutes of Health awarded to JDL.

## Declaration of competing interest

The authors declare no conflicts of interest.

## Data availability

Data will be made available upon reasonable request from the corresponding author.

## Notes

### Competing Interest Statement

The authors have declared no competing interest.

